# Overexpression of α–synuclein in Midbrain Dopamine Neurons Reduces Dopamine Release Without Cell Loss and Drives Mild Motor Deficits in Mice

**DOI:** 10.1101/2025.03.17.643840

**Authors:** Kelsey Barcomb, Jen-Tsiang T. Hsiao, Julia Lemak, Xiaowen Zhuang, Berenice Coutant, Anthony Yanez, Aditya Behal, Yuhong Fu, Talia Lerner, Ken Nakamura, Robert Edwards, Zayd Khaliq, Glenda Halliday, Chris Ford, Alexandra B. Nelson

**Affiliations:** Department of Pharmacology, University of Colorado, Anschutz, Denver, CO; University of Sydney, Sydney, Australia; Department of Neurology, UCSF, San Francisco, CA 94158, USA; National Institute of Neurological Disease and Stroke, Bethesda, ML, USA; Department of Physiology, Northwestern University, Chicago, IL, USA; Gladstone Institute of Neurological Disease, Gladstone Institutes, San Francisco, CA 94158, USA; Kavli Institute for Fundamental Neuroscience, UCSF, San Francisco, CA 94158, USA; Weill Institute for Neurosciences, UCSF, San Francisco, CA 94158, USA; Aligning Science Across Parkinson’s (ASAP) Collaborative Research Network, Chevy Chase, MD 20815, USA

## Abstract

It has proven challenging to faithfully recapitulate the key pathological, physiological, and behavioral features of Parkinson’s Disease (PD) in animals. Here we used adeno-associated virus (AAV) vectors to achieve cell type-specific overexpression of wild-type human *α*-synuclein (*α*syn) and a fluorophore (mCherry) in midbrain dopamine neurons to model PD in mice. We found that AAVs drove selective expression of both *α*syn and mCherry in midbrain dopamine neurons. In conjunction with approximately 2-fold overexpression of *α*syn, we found several histopathological markers of PD-like pathology, including progressive accumulation of phosphorylated and aggregated *α*syn, ubiquitin, and a reduction in the expression of tyrosine hydroxylase, without overt cell loss. In parallel, *α*syn overexpression drove a profound loss of evoked dopamine release, without a substantive change in the intrinsic properties of dopamine neurons, nor in striatal dopamine content. Finally, *α*syn overexpression led to mild locomotor deficits. Together, these findings suggest that moderate *α*syn overexpression can mimic some aspects of premotor and early symptomatic phases of PD, including markers of Lewy Body-like pathology and functional loss of evoked dopamine release. This model may be useful for investigating cellular and circuit mechanisms related to PD pathogenesis and progression.

## Introduction

Parkinson’s Disease (PD) is a neurodegenerative disease characterized by progressive motor, cognitive, and behavioral symptoms. While the onset, rate of progression and symptom severity vary between individuals, the unifying pathological feature of PD is the accumulation of *α*synuclein (*α*syn) in neurons^1,2^. In people with PD, pathological *α*syn is found in many brain areas, but is especially prominent in midbrain dopamine neurons of the substantia nigra, pars compacta (SNc). Substantial dopamine cell loss in the SNc, and axonal loss in its major synaptic target, the striatum, is closely correlated with the onset and severity of classical motor symptoms of PD^3-6^.

In healthy individuals, *α*syn is expressed throughout the brain, and is enriched at synapses^7^. Although the normal function of *α*syn is not entirely clear, several groups have linked *α*syn to the release and/or reuptake of synaptic vesicles^7,8^. In rare cases of PD, duplications or triplications of the *α*syn gene, SNCA, have been identified^9,10^, suggesting that overexpression of *α*syn may be causally related to the development of PD. These findings suggest that excess and/or pathologically accumulated *α*syn may drive alterations in cellular function, including synaptic release.

Animal models are a tool to study the pathogenesis and pathophysiology of PD, particularly how it begins and progresses. Ideally, such models would show (1) typical PD brain pathology, including pathological *α*syn or other neurodegenerative changes, (2) typical PD signs/symptoms, and (3) gradual progression. It has been challenging to create a model that fulfills all of the above criteria. Toxin-based models recapitulate dopamine cell loss and several of the typical motor symptoms of PD^11^, but do not typically show *α*syn pathology and are not progressive in nature. Transgenic models based on human PD genes often show neither classic pathology nor frank motor symptoms. Rodent models based on the overexpression of either wild-type or mutant *α*syn have been developed over the past two decades^12-14^. These models are variable, but in many cases show progressive *α*syn pathology and in some cases cell/axonal loss and/or progressive motor deficits. Whether using transgenic or viral methods, most of these models do not specifically target SNc dopamine neurons, instead using ubiquitous promoters to drive widespread overexpression^12,15^ or viral injection to target *α*syn to the midbrain, but not in dopamine neurons specifically.

Here, we use a cell-type specific viral strategy to drive overexpression of wild-type human *α*–syn in the midbrain dopamine neurons of mice. In this model, we find little evidence of midbrain cell loss, but accumulation of pathological *α*syn, progressive Lewy Body-like pathology, and reductions in a key enzyme in dopamine metabolism, tyrosine hydroxylase. In parallel with this pathology, we observe changes in dopamine neuron function, including evoked dopamine release. Finally, animals overexpressing *α*syn show progressive motor deficits, which correlate with the development of Lewy Body-like pathology. This model may be especially useful in investigating the factors governing onset and progression of PD, and the contribution of altered cellular functions, such as synaptic release, versus frank cell loss.

## Results

To achieve cell type-specific overexpression of wild-type human *α*syn in midbrain dopamine neurons, we combined a Cre recombinase mouse line targeting dopamine neurons (DAT-IRES-Cre^16^) and bilateral injection of AAV encoding Cre-dependent *α*syn and mCherry or mCherry alone (Figure 1A). Fusing proteins with fluorophores can sometimes create large proteins that impede transport and targeting^17^, so we separated *α*syn and mCherry expression with an internal ribosomal entry sequence (IRES) in the AAV. DAT-Cre mice were randomized and interleaved into *α*syn-mCherry or mCherry groups.

**Figure 1:**
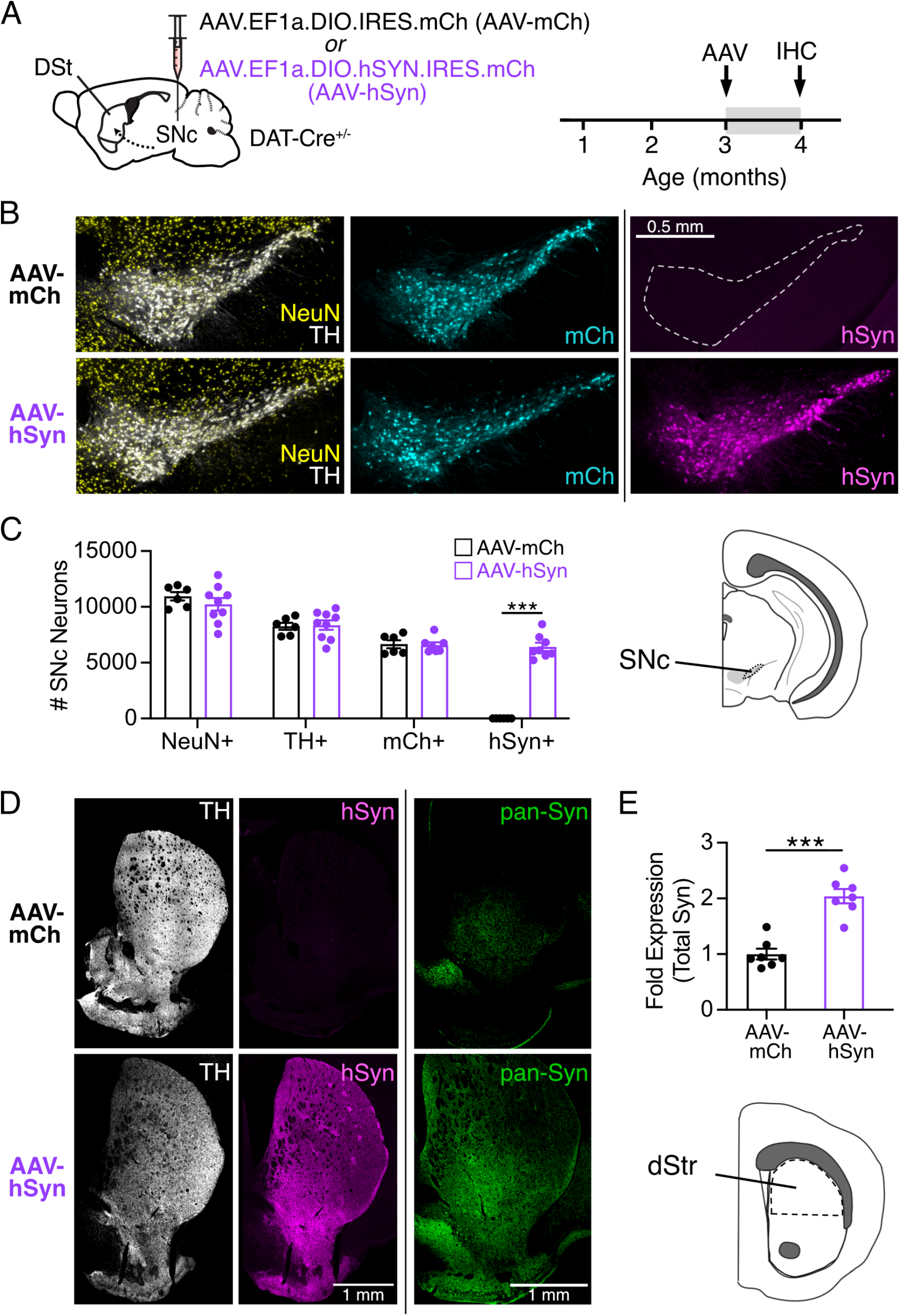
AAV-based overexpression of wild-type human *α*-synuclein in midbrain dopamine neurons. DAT-Cre mice were injected in the substantia nigra pars compacta (SNc) with AAV encoding mCherry (AAV-mCh) or human wild-type *α*-synuclein and mCherry (AAV-hSyn), and sacrificed after 1 month for postmortem histology. (A) Experimental timeline. (B) Representative midbrain sections from mice injected with AAV-mCh (top) or AAV-hSyn (bottom). From left to right, images show all neurons (NeuN, yellow) and dopamine neurons (tyrosine hydroxylase, TH white), mCherry (cyan) and human *α*-synuclein (hSyn, magenta). hSyn immunostain image is from a different brain section, as indicated by the vertical line separating the images. (C) Summary of SNc neurons expressing NeuN, TH, mCherry, and human *α*-synuclein in each group. There was no difference in the expression of mCherry between the two viruses, but only AAV-hSyn injected animals expressed human α-synuclein, p < 0.001. (D) Representative striatal sections from mice injected with AAV-mCh (top) or AAV-hSyn (bottom). From left to right, images show TH-expressing fibers (TH, white), human *α*--synuclein (hSyn, magenta), and total *α*-synuclein (pan-Syn, green). (E) Summary of the fold-overexpression of *α*-synuclein, calculated as the fluorescence intensity of total *α*-synuclein immunohistochemistry in the dorsal striatum over the average *α*-synuclein in the AAV-mCh (control group). In the AAV-hSyn group, expression was approximately 2-fold higher than in the control group (p < 0.0001). AAV-mCh: N=6-7; AAV-hSyn: N=6-9. All data shown as mean +/-SEM. Superimposed dots represent individual mice.

We first tested whether our two AAVs were expressed in SNc dopamine neurons, using postmortem histology 1 month after viral injection. Animals injected with the undiluted AAVs encoding mCherry alone or *α*syn-mCherry showed robust expression of mCherry in SNc cell bodies expressing the neuronal marker NeuN and tyrosine hydroxylase (TH), a marker of dopamine neurons (Figure 1B). In the *α*syn-mCherry group, there was also robust expression of human *α*–syn as assessed with the human-specific *α*–syn antibody 15G7^18^. Overall, undiluted AAV achieved good coverage of the SNc: the majority of dopamine neurons expressed mCherry in both groups, and human *α*–syn in the *α*–syn-mCherry group (Figure 1C). These findings suggest our AAVs drove robust and selective expression of *α*–syn and/or mCherry in midbrain dopamine neurons.

The axons of dopamine neurons are a major site for pathology in PD. To test whether our AAVs drove strong expression in the striatum, the primary synaptic target of dopamine neurons, we performed immunohistochemistry in the striatum. We observed expression of mCherry in the axon terminals of dopamine neurons within the striatum in both groups, but found human *α*–syn only in the *α*–syn-mCherry group (Figure 1D). The degree of *α*–syn overexpression may correlate with the rapidity of disease progression in PD^19^. To assess the degree of overexpression in our AAV-injected mice, we used immunohistochemistry for total *α*–syn (mouse and human, with the pan-synuclein antibody syn-1), and compared *α*–syn-mCherry and mCherry groups. *α*–syn expression in the dorsal striatum of *α*–syn -mCherry mice was just over 2-fold that seen in mCherry controls (Figure 1E). These data validate that our AAV drives strong expression of human *α*–syn in both the cell bodies and axons of midbrain dopamine neurons.

PD is a progressive disease, in terms of both pathology and behavioral manifestations. We next tested whether AAV-mediated *α*–syn overexpression in midbrain dopamine neurons produced progressive histopathology at 1, 2, and 4 months after AAV injection (Figure 2A). We first assessed the expression of TH, as a marker for dopamine neurons and the most common readout of PD-like pathology in animal models. In the striatum, we found that the intensity of TH immunoreactivity was reduced at all timepoints in the *α*–syn-mCherry group (Figure 2B-C). The reduction in TH expression could be associated with a loss of dopamine neurons, dopamine axons, or a functional change in dopamine neurons. To differentiate between these possibilities, we examined the integrity of dopamine neurons in the midbrain. The number of neurons (labeled with NeuN) in the SNc did not decline in the mCherry control group, nor in the *α*syn-mCherry group (Figure 2D-E). As the vast majority of SNc neurons are dopamine neurons, this observation suggests that there is not a significant reduction in the number of dopamine neurons in the *α*–syn-mCherry group. At 2 months, however, the fraction of SNc neurons staining positive for TH, was lower in the *α*–syn-mCherry group (though this was not significant as compared to the control group at the same timepoint) (Figure 2F). These findings indicate that overexpression of *α*–syn leads to a reduction in striatal TH expression, without overt loss of cell bodies, which may suggest a change in the function of midbrain dopamine neurons.

**Figure 2:**
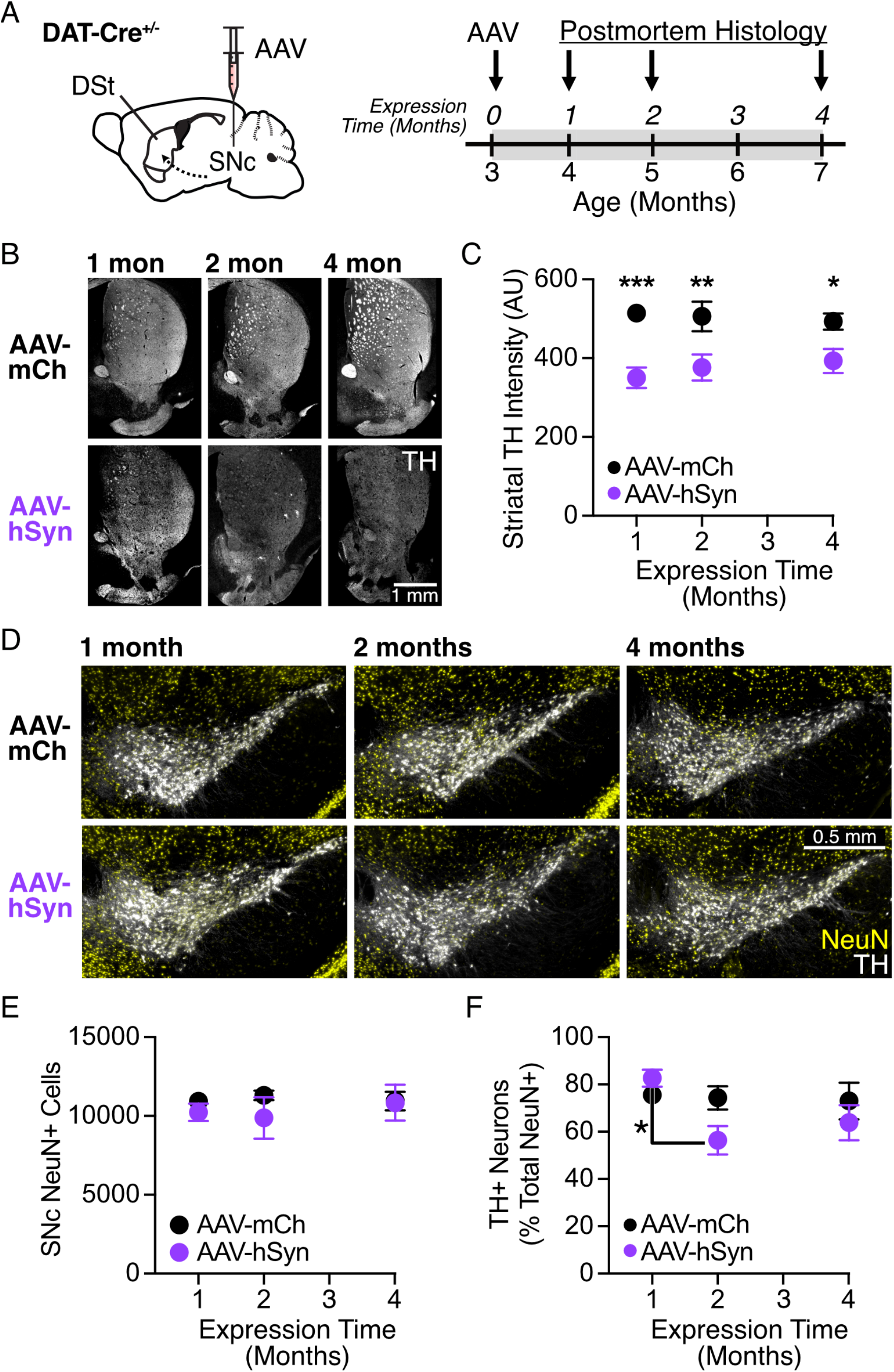
Overexpression of *α*-synuclein reduces TH expression, but does not cause loss of midbrain dopamine neurons. DAT-Cre mice were injected in the SNc with AAV encoding mCherry or human wild-type *α*-synuclein and mCherry, and sacrificed after 1, 2, or 4 months for postmortem histology. (A) Experimental timeline. (B) Representative striatal sections from mice injected with AAV-mCh (top) or AAV-hSyn (bottom). Tyrosine hydroxylase (TH) staining is shown in mice sacrificed at 1, 2, and 4 months after AAV injection. (C) Average striatal TH immunofluorescence intensity was decreased at all timepoints in mice injected with AAV-hSyn. (p < 0.0001, Post Hoc Tukey’s Test, p < 0.0002, p = 0.001, p = 0.02, for 1m, 2m, and 4m respectively) (D) Representative midbrain sections from mice injected with AAV-mCh (top) or AAV-hSyn (bottom), stained for NeuN (yellow) and TH (white) after 1, 2 or 4 months. (E) Summary of the total number of neurons (NeuN-positive) in the SNc. There was no significant change between groups or across timepoints (p = 0.25). (F) Summary of TH-positive neurons, as a percentage of the total number of neurons (NeuN) in the SNc. In the hSyn group, there was a reduction in the percentage of TH-positive neurons at 2 versus 1 month, but no other significant differences within or between groups were detected (p = 0.04, Post Hoc Tukey’s, p = 0.02). AAV-mCh: N = 6, 6, 6 at 1, 2, and 4 months. AAV-hSyn: N = 9, 5, 7 at 1, 2, and 4 months. All data shown as mean +/-SEM.

As PD progresses, *α*syn aggregates, leading to a number of histopathological changes in midbrain dopamine neurons and their processes, including the eventual formation of Lewy neurites and Lewy bodies^1^. We examined our postmortem tissue for some of these histopathological hallmarks across the 1, 2, and 4 month timepoints. Between 1 and 2 months, we observed an increase in beaded processes in *α*–syn-expressing dopamine neurons within the SNc (Figure 3A). Lewy bodies are complex structures, containing aggregated, fibrillar *α*–syn, as well as ubiquitin. To assay for Lewy body-like pathology in our AAV model, we stained the SNc for phosphorylated *α*syn, p62, and ubiquitin (Figure 3B-C), and the striatum for human fibrillar *α*syn (Figure 3D). Between 1 and 2 months after AAV injection, we detected the emergence of phospho-Ser129 *α*syn in SNc dopamine neurons, which persisted through 4 months (Figure 3B). Over time, we saw a gradual increase in the ubiquitin binding protein p62, as well as Thioflavin S, a marker of fibrillar protein aggregates, in SNc dopamine neurons (Figure 3C). Finally, we examined the axonal arbors of dopamine neurons for PD-like pathology. In the striatum, we found an increase in aggregated *α*syn, as measured by the human disease-specific antibody 5G4^20^. Together, these observations suggest that AAV-mediated overexpression of wild-type human *α*–syn can drive many of the early histopathological hallmarks of PD, though without overt cell loss.

**Figure 3:**
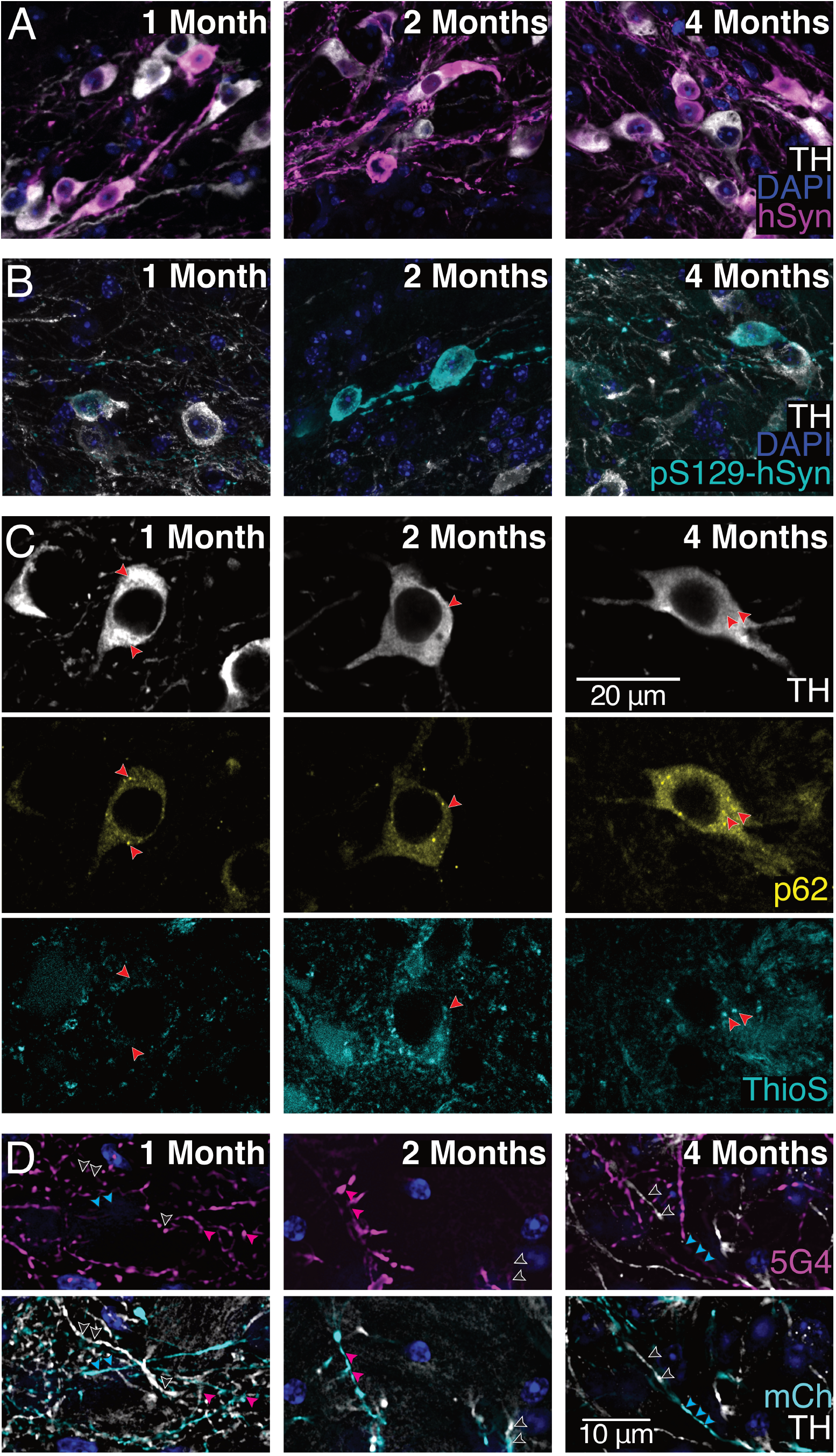
Synuclein overexpression leads to Lewy Body-like pathology in midbrain dopamine neurons. High magnification images of SNc cell bodies (A-C) and striatal axons (D) at 1, 2 and 4 months from mice injected with AAV-hSyn. (A) Tyrosine hydroxylase (TH, white), DAPI (blue) and human *α*-synuclein (hSyn, magenta) staining shows a reduction in TH immunoreactivity and more extensive beading of processes at 2 months. (B) TH (white), DAPI (blue) and pS129-hSyn (cyan) staining showing pS129 immunoreactivity at 2 and 4 months. (C) TH (white), p62 (yellow) and thioflavin S (ThioS, cyan) staining shows pS129 and p62 immunoreactivity at 2 and 4 months. (D) Neurons containing TH (white) and p62 (cyan) immunoreactivity and thioflavin-S (thioS, pink) fibril fluorescence at 2 and 4 months. (D) Immunohistochemistry for TH (white), DAPI (blue), mCherry (cyan), and aggregated *α*-synuclein (5G4, pink) in striatal sections at 1, 2 and 4 months shows a reduction in TH-positive axons at 2 and 4 months, which parallels the development of fibrillar *α*-synuclein pathology. Arrowheads demarcate axonal regions for comparison across sections, and highlight the loss of TH in regions of the axon that show aggregated *α*-synuclein pathology.

In PD and several animal models of PD, neurodegenerative loss of midbrain dopamine neurons leads to reduced levels of dopamine and dopamine release. Many hypothesize that before cell loss, dopamine neurons may be dysfunctional, with either increased or decreased activity^21^. Given the interactions between *α*syn and membranes, including synaptic vesicles^7^, we hypothesized that AAV-mediated *α*syn overexpression would alter synthesis and/or synaptic release of dopamine. To test this hypothesis, we measured evoked dopamine release in *ex vivo* brain slices. We crossed DAT-Cre mice to the Cre-dependent Channelrhodopsin (ChR2) line, Ai32, to achieve expression of ChR2 in dopamine neurons, then injected AAVs encoding *α*syn and mCherry or mCherry alone (Figure 4A). After 1 month of AAV expression, we prepared acute striatal slices for fast-scan cyclic voltammetry (FSCV; Figure 4B, top). Using a carbon fiber microelectrode, we recorded optically-evoked dopamine release (Figure 4B, bottom). In the dorsomedial striatum (DMS) and dorsolateral striatum (DLS) of control mice, we observed robust dopamine release in response to a single blue light flash (1 msec, 470 nM) (Figure 4C). However, in slices from *α*–syn-mCherry-expressing mice, evoked release was markedly reduced (Figure 4C). Even after only 1 month of AAV expression, evoked dopamine release was less than half that seen in control slices. A reduction in dopamine release may be mediated by loss of dopamine axons, reduced dopamine synthesis in intact axons, or an alteration in the functional properties of dopamine neurons. To distinguish between these possibilities, we first tested striatal dopamine content. Given the reduction in the expression of TH, the rate limiting enzyme for the synthesis of dopamine, in *α*syn-mCherry mice, we hypothesized the striatal dopamine content might be reduced in these animals. Surprisingly, we found intact tissue dopamine content in striatal tissue samples from *α*syn-mCherry mice vs mCherry controls (Figure 4D). These findings suggest that the reduction in evoked dopamine release caused by overexpression of *α*syn in midbrain dopamine neurons might result from a change in the functional properties of dopamine neurons.

**Figure 4:**
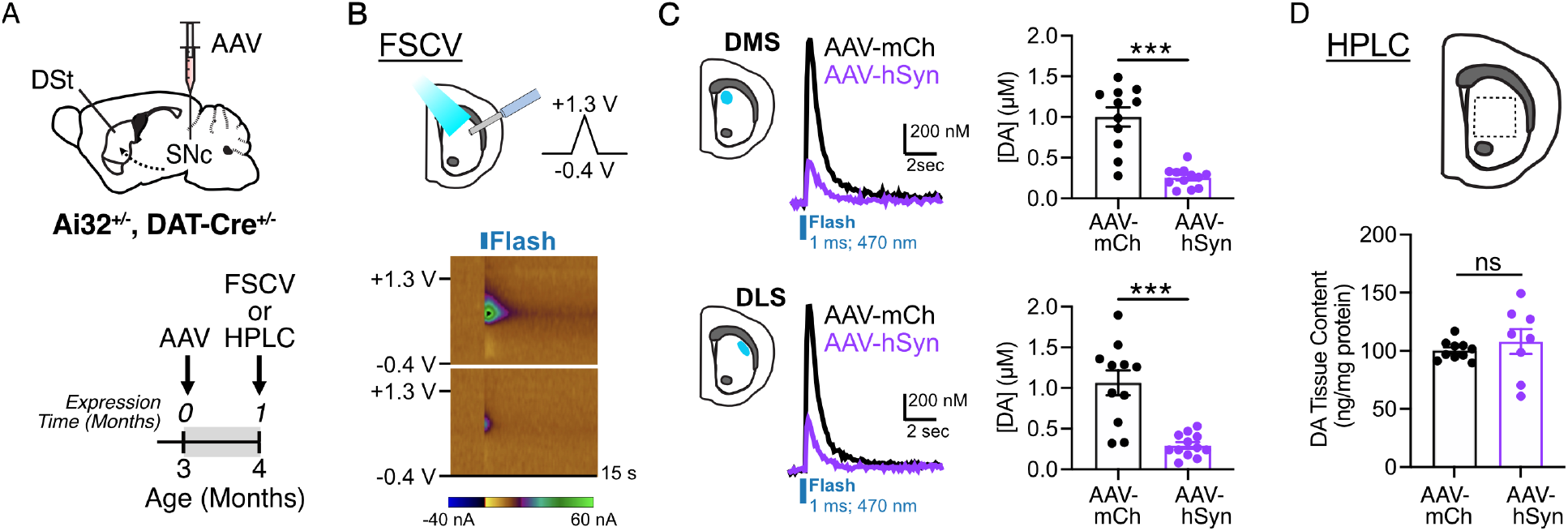
Synuclein overexpression leads to progressive loss of evoked dopamine release. DAT-Cre;Ai32 mice were injected in the SNc with AAV encoding mCherry or human wild-type *α*-synuclein and mCherry, and sacrificed after 1 month for measurement of evoked dopamine release (A-C) or assessment of tissue dopamine content (D). (A) Experimental timeline. (B) Cartoon (top) of ex vivo fast-scan cyclic voltammetry (FSCV) approach using blue light pulses (1 ms, 470 nm) to evoke dopamine release from ChR2-expressing dopamine terminals in the striatum. The triangular voltage input yields current responses, shown as heatmaps from mice injected with AAV-mCh or AAV-hSyn. Dopamine release is shown over a 15 second sweep, with a green color indicating higher dopamine release. (C) Representative FSCV trace showing calculated dopamine release (in nM) in recordings from the dorsomedial striatum (DMS; top) and dorsolateral striatum (DLS; bottom) of AAV-mCh and AAV-hSyn injected mice. At right are summaries of evoked dopamine release across groups. DA release was significantly reduced in slices from AAV-hSyn injected mice in both the DMS (p < 0.0001) and DLS (p = 0.0002) AAV-mCh: N = 11; AAV-hSyn: N = 12. (D) Cartoon showing area for striatal tissue punch (top), and High-Performance Liquid Chromatography (HPLC) measurement of tissue dopamine content across AAV-mCh and AAV-hSyn mice (bottom). Tissue dopamine was not different between AAV-mCh and AAV-hSyn injected mice (p = 0.27). AAV-mCh: N = 10; AAV-hSyn: N = 8. All data shown as mean +/-SEM. Superimposed dots represent individual mice.

Evoked dopamine release may be regulated by the intrinsic and synaptic properties of dopamine neurons. To determine whether *α*–syn overexpression altered the intrinsic properties of dopamine neurons, we performed ex vivo patch-clamp recordings of SNc dopamine neurons. Given the profound reductions in evoked dopamine release after only 1 month of AAV expression, we examined intrinsic properties at the same time point, targeting mCherry-positive SNc dopamine neurons (Figure 5A). Dopamine neurons are tonically active *in vivo* and in *ex vivo* slice preparations, and this tonic firing is one readout of their intrinsic excitability. We found no difference in the spontaneous firing rate of SNc dopamine neurons in mCherry vs *α*syn-mCherry mice (Figure 5B). Likewise, we did not observe a difference in the resting membrane potential (Figure 5C) or several other intrinsic properties (Supplementary Table 2). To further probe the intrinsic excitability of SNc dopamine neurons, we injected small pulses of current to evoke firing, and constructed an input current-output firing rate (IF) plot (Figure 5D). Using this readout, *α*–syn-expressing dopamine neurons actually showed increased excitability (Figure 5D, right). We next examined the spontaneous action potential waveform, finding modest changes, including a small reduction in the action potential half-width (Figure 5E) accompanied by a decrease in the downstroke velocity and increase in the upstroke velocity (Table 2). These recordings of SNc dopamine neurons indicate that overexpression of *α*syn does not cause marked changes in intrinsic properties, and those that are observed would not explain the profound reduction in evoked dopamine release seen with fast-scan cyclic voltammetry.

**Figure 5:**
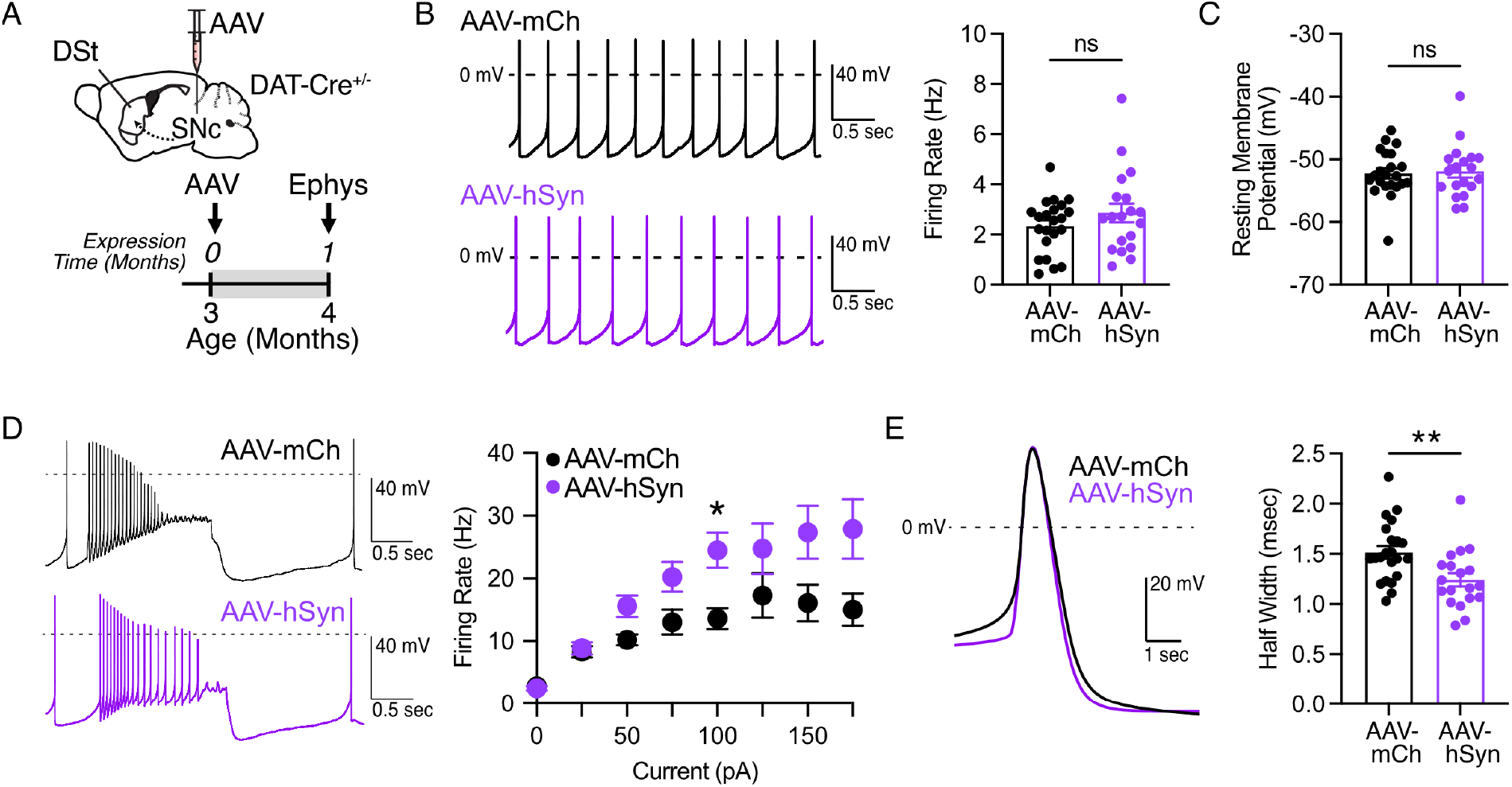
Synuclein overexpression does not alter the intrinsic properties of SNc dopamine neurons. DAT-Cre mice were injected in the SNc with AAV encoding mCherry or human wild-type *α*-synuclein and mCherry, and sacrificed after 1 month for *ex vivo* patch-clamp recordings of SNc dopamine neurons. (A) Experimental timeline. (B) Representative whole-cell recordings from mCherry-expressing SNc neurons from the AAV-mCh (top) and AAV-hSyn (bottom) injected groups. Summary of spontaneous firing rates is shown at right. No difference was observed between the groups (p = 0.23). (C) Summary of resting membrane potential. No difference was observed between the groups (p = 0.80). (D) Representative whole-cell recordings showing the response to a square wave current injection. Summary of input-output curve is shown at right. (p < 0.0001, Post Hoc Bonferroni’s Test, p = 0.02). (E) Average spike shape in representative neurons from the AAV-mCh and AAV-hSyn groups (left) and summary of the spike half-widths (right). The half-width was significantly reduced in the AAV-hSyn group (p = 0.005). AAV-mCh: N=5, n=22; AAV-hSyn: N=6, n=19. N=animals; n=cells. All data shown as mean +/-SEM. Superimposed dots represent individual cells.

The clinical hallmark of PD is progressive motor dysfunction, including bradykinesia, rigidity, tremor, and postural instability. Based on the neuropathological and functional changes we observed, without overt cell loss, we hypothesized *α*syn-overexpressing mice would show mild motor deficits. To test this hypothesis, we performed longitudinal, blinded behavioral assessments at three timepoints after AAV injection: 1, 2 and 4 months (Figure 6A). At each timepoint, we used three common assays of motor function in mice: the open field, accelerating rotarod, and pole tests. The open field test measures locomotor velocity, which is reduced in several mouse models, particularly those with overt loss of midbrain dopamine neurons. We found no differences between mCherry and *α*syn-mCherry groups at the earliest time point (1 month), but reductions in movement velocity in the *α*syn-mCherry groups at 2 months (Figure 6B). The accelerating rotarod test assays several aspects of motor function, and performance is impaired in several mouse models of PD^12,22^. As the rod turns at higher frequencies (up to 80 rpm), maintaining balance becomes more difficult, and eventually even healthy mice fall off the rod. Using the time to fall off the rotarod as the primary outcome measure, we did not find impairments in *α*syn-overexpressing mice (Figure 6C). Finally, we performed the pole test. The pole test assays how quickly a mouse, placed facing upwards on a narrow vertical pole, is able to re-orient (facing downwards) and descend to the base. The pole test is hypothesized to reflect both fine and gross motor coordination, and is also impaired in some mouse models of PD^12,23^. We found a high level of variance in performance on the pole test, but no significant differences between *α*syn-mCherry and mCherry mice (Figure 6D). These data show that *α*–syn-overexpressing mice have mild locomotor deficits in the open field, without clearly impaired performance in the accelerating rotarod or pole test.

**Figure 6:**
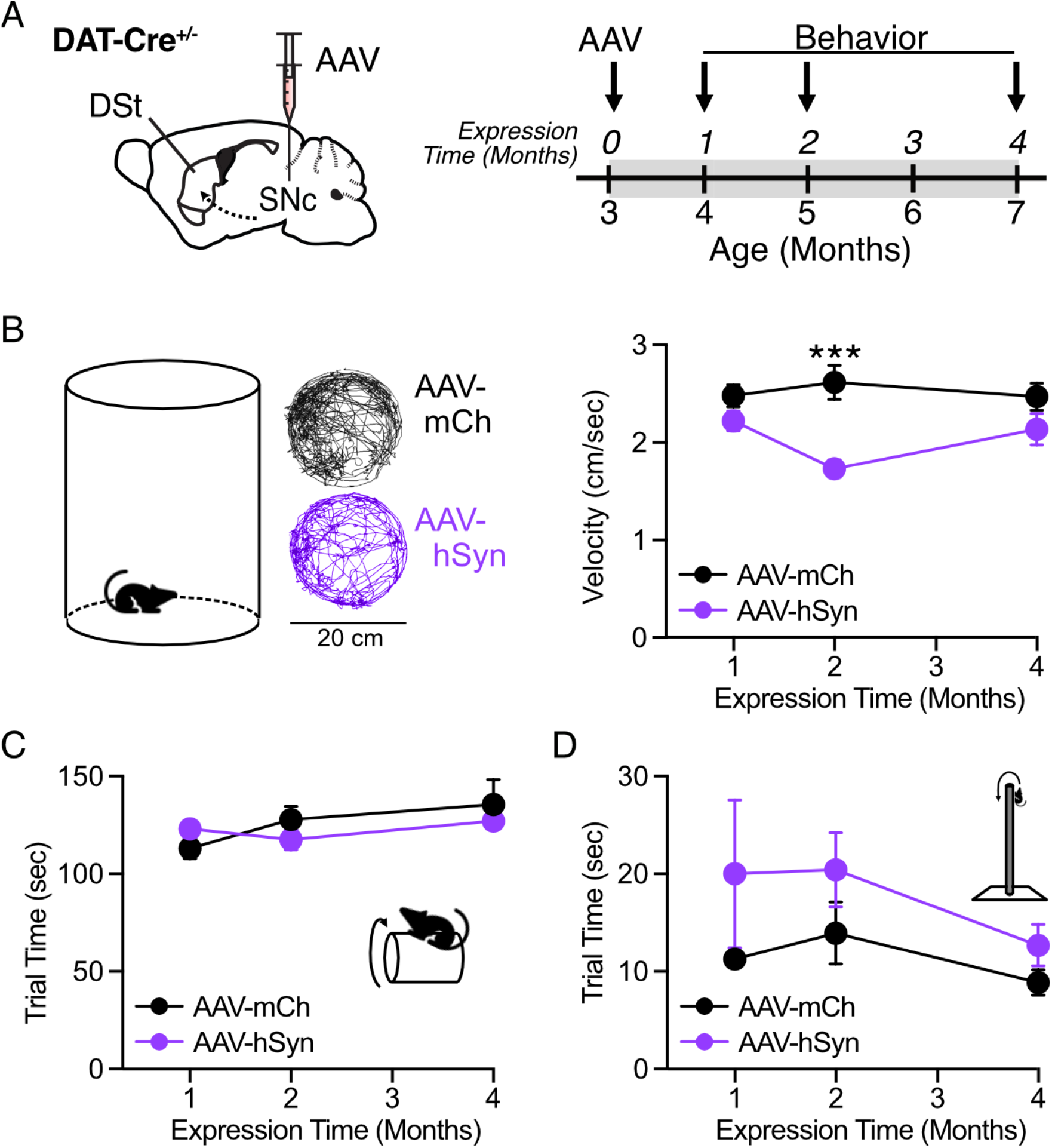
Synuclein overexpression in midbrain dopamine neurons leads to motor deficits. DAT-Cre mice were injected in the SNc with AAV encoding mCherry or human wild-type *α*-synuclein and mCherry. Motor assays were performed at 1, 2, and 4 months. Mice were sacrificed for postmortem histology and/or physiology at one of the three timepoints, so not all animals were recorded across all three timepoints. (A) Experimental Timeline. (B) Reduced open field movement in mice overexpressing *α*-synuclein. Left: Representative traces of movement in the open field, extracted from overhead video tracking, in AAV-mCh and AAV-hSyn groups. The open field test had a duration of 20 minutes in a cylinder of 20 cm diameter. Right: Summary of average locomotor velocity in the AAV-mCh and AAV-hSyn groups. Velocity was reduced at 2 months in the AAV-hSyn group, as compared to the control group (p = 0.03, Post Hoc Bonferroni’s Test, p < 0.0001). (C) Performance on the accelerating rotarod test was comparable between groups. Data represent the best trial performance (longest time on rotarod before a fall) across three trials for each mouse at each time point. (D) Performance on the pole test was comparable between groups. Data represent the best trial performance (fastest descent of pole) across three trials for each mouse at each time point. . AAV-mCh: N = 31, 23-24, 12 across 1, 2, and 4 months. AAV-hSyn: N = 31, 22, 15 across 1, 2, and 4 months. There were fewer completions of the pole test at 1 month, where N = 22 and 27 for AAV-mCh and AAV-hSyn. All data shown as mean +/-SEM.

In the previous experiments, we used undiluted AAVs. Though we did not find significant direct toxicity of the mCherry or *α*syn-mCherry AAVs, we were interested in whether diluting the AAV could produce similar histopathological, physiological, and/or behavioral effects. We were particularly interested in whether there were different thresholds for inducing physiological versus histopathological changes. We diluted each AAV in a 1:4 ratio with sterile saline (1 uL of AAV to 4 uL of saline), and injected the same volume at the same location in the bilateral SNc of DAT-Cre mice. The diluted viruses yielded slightly lower coverage of TH+ SNc neurons, though still more than half of neurons showed mCherry expression at 1 month (Supplementary Figure 1A, left). Like the undiluted AAV, the diluted AAV encoding *α*syn-mCherry yielded robust synuclein expression, but did not cause overt cell loss, assayed with NeuN at any of the three timepoints (Supplementary Figure 1A, middle). The diluted AAV did not cause a reduction in the number of TH+ SNc neurons at any time point (Supplementary Figure 1A, right). To determine whether the diluted AAV disrupted dopamine signaling, we performed biochemical and physiological assays after 1 month of expression. As with the undiluted virus, we did not observe a decrease in striatal dopamine content in the *α*syn-mCherry group (Supplementary Figure 1B). However, the diluted virus did lead to a robust decrease in evoked dopamine release, as measured by fast-scan cyclic voltammetry, in both the DMS and DLS (Supplementary Figure 1C). Finally, we performed the same blinded motor behavior assays (open field, accelerating rotarod, and pole tests) across 1, 2, and 4 months after AAV injection. Animals in the diluted mCherry and *α*syn-mCherry groups showed qualitatively similar performance to those in the undiluted groups, with a reduction in open field velocity seen at 2 months in the *α*syn-mCherry group, but no significant differences in the accelerating rotarod or pole test (Supplementary Figure 1D-F; Figure 6). Together, these results with diluted AAV suggest that there is a lower threshold for *α*syn overexpression-mediated reductions in evoked dopamine release and mild locomotor deficits, as compared to reductions in TH expression.

## Discussion

We found that AAV-mediated cell type-specific wild-type human *α*–syn overexpression in midbrain dopamine neurons leads to progressive nigrostriatal histopathology, physiological dysfunction, and mild motor deficits in mice. Moderate (2-fold) overexpression led to a reduction in TH expression, and a progressive increase in markers of *α*–syn aggregation, such as phosphorylated *α*–syn, p62, and thioflavin S. We found this was paralleled by a profound loss of evoked dopamine release, despite the fact that there was neither overt cell loss in the SNc, nor a reduction in striatal dopamine content. Finally, we found that at 2 months of expression, mild locomotor deficits developed.

Our approach shares features with several *α*syn overexpression models, including both transgenic and viral-based rodent models. BAC transgenic mice broadly overexpressing human *α*syn, approximately 2-fold in the striatum (“ SNCA-OVX” mice), show many parallels. After a longer period of overexpression (18 months), SNCA-OVX mice show a small reduction in the number of midbrain TH+ cells (and overall number of cells), as well as deficits on the rotarod and pole test^12^. These mice also show a modest reduction in evoked dopamine release, without a decrease in dopamine content. Transgenic rats expressing human *α*syn using a similar approach also have decreased TH+ cells and mild motor deficits at 12-13 months, which are paralleled by an increase in dopamine neuron excitability^24,25^. Compared to these transgenic models, prior viral *α*syn overexpression models typically show more rapid induction of cellular toxicity, loss of TH expression, and behavioral deficits. A common caveat in both transgenic and viral *α*syn models is the broad overexpression of *α*syn. With viral models, *α*syn overexpression can be directed to a specific region, commonly the midbrain, but is not specific to a cell type, such as SNc dopamine neurons, that is particularly vulnerable to PD pathology. Another caveat is the potential for toxicity by either the virus itself or fused fluorophores. Indeed, in some studies, a construct expressing *α*syn alone is compared to a construct expressing GFP alone^13^. To avoid these limitations, we created two constructs, one control and one for *α*syn overexpression, which were identical in every way except for the *α*syn. Moreover, we used an IRES between *α*syn and fluorophore elements, to promote expression of both proteins and reduce toxicity.

We found several indications of axon-first pathology in the *α*syn overexpression model. In the striatum, TH was downregulated, even at the earliest timepoint (1 month), while there was no cell body loss at any timepoint, and significant *α*syn pathology developed in cell bodies only at later timepoints (2, 4 months), as assayed with traditional markers like phosphorylated *α*syn, p62, and Thioflavin S. The profound loss of evoked dopamine release was also noted at the earliest timepoint (1 mo), consistent with early synuclein-mediated synaptic impairment. These observations are consistent with prior reports of early defects in dopamine release, even without cell loss, in both PD and animal models of PD^21^. While we observed alterations in dopamine axonal morphology with *α*syn overexpression, we were unable to determine if there was a loss of axonal integrity. The normal dopamine content, however, suggests that presynaptic terminals are either intact or dopamine content is upregulated in surviving terminals.

We found a significant reduction in evoked dopamine release in the *α*syn overexpression model, without differences in dopamine content, nor marked changes in the intrinsic properties of dopamine neurons, at the cell body level. This observation is in line with findings that altering *α*syn expression level regulates synaptic properties^26,27^, potentially via synuclein’s interaction with membranes, including the synaptic vesicle. Notably, amongst the different pathological, physiological and behavioral outcomes measured here, evoked dopamine release was the first to emerge, even in animals treated with diluted AAV. This observation is also consistent with the idea of physiological dysfunction prior to overt pathology.

Parkinson’s Disease is characterized by classical motor signs and symptoms, such as rigidity, slowing of movement, alterations in gait and balance. Some of these features can be recapitulated in animal models. In our cell type-specific AAV model of *α*syn overexpression, we observed very mild motor deficits, primarily manifest as a slowed locomotor activity in the open field test. We did not observe parallel changes in performance on the accelerating rotarod and pole test. While these tests are commonly used in animal models of PD, and are often reported to be more sensitive for early motor phenotypes, we found that there was relatively high variance in these measures, and with the sample sizes used here, we did not detect a difference between control animals and those overexpressing *α*syn. The modest motor phenotype may reflect that there is not significant SNc cell loss. Though evoked dopamine release was reduced in the model, this decrease may be below the threshold for driving classical motor signs of PD. We did not test these animals for other features of PD, but striatal dopamine release has been linked to functions such as motor learning, cognitive flexibility, and sleep^28,29^.

AAV-mediated overexpression of *α*syn drove sequential changes in in synaptic transmission, expression of cell markers such as TH, followed by the development of *α*syn histopathology and mild motor deficits. This model thus may be useful to study the premotor and/or prodromal features of PD, as well as the progression of disease. The cell type-specific nature of the model will also allow future testing of the specific role of *α*syn and *α*syn-mediated alterations in synaptic function in different PD-associated brain areas and cell types.

## Acknowledgments

This research was funded in whole or in part by Aligning Science Across Parkinson’s [ASAP020529] through the Michael J. Fox Foundation for Parkinson’s Research (MJFF). For the purpose of open access, the author has applied a CC BY public copyright license to all Author Accepted Manuscripts arising from this submission. We also thank Vivienne Yuan and Amy Burke for their technical support on histopathology work, and Kohei Kano for his help in validating synuclein expression levels in the AAV model.

