## Supplemental Figures, Tables, and Methods for "Overexpression of α–synuclein in Midbrain Dopamine Neurons Reduces Dopamine Release Without Cell Loss and Drives Mild Motor Deficits in Mice"

### Supplementary Material

**Fig 1C – Cell counts in SNc, AAV-mCh v hSyn**

| | AAV-mCh | | | AAV-hSyn | | | |
|---|---|---|---|---|---|---|---|
| | Mean | SEM | n | Mean | SEM | n | |
| NeuN+ (# cells) | 10948.5 | 381.4 | 6 | 10230.8 | 555.0 | 9 | ns |
| TH+ (# cells) | 8272.7 | 314.0 | 6 | 8371.8 | 412.5 | 9 | ns |
| mCh+ (# cells) | 6660.0 | 348.6 | 6 | 6573.7 | 261.6 | 7 | ns |
| hSyn+ (# cells) | 0.0 | 0.0 | 6 | 6430.3 | 368.2 | 8 | *** |

2 Way ANOVA
Main Effect of Cellular Marker: $F(3,49) = 118.6$, ***$p < 0.0001$
Main Effect of Virus: $F(1,49) = 24.86$,, ***$p < 0.0001$
Interaction: $F(3,49) = 33.95$, ***$p < 0.0001$

Bonferroni's Post Hoc Test:
hSyn+: AAV-mCh < AAV-hSyn, ***$p < 0.0001$

**Fig 1E – Fold overexpression of a-synuclein in SNc**

| | AAV-mCh | | | AAV-hSyn | | | |
|---|---|---|---|---|---|---|---|
| | Mean | SEM | n | Mean | SEM | n | |
| Pan-Syn IF / Total | 1.00 | 0.10 | 7 | 2.04 | 0.13 | 7 | *** |

Unpaired t-test
$t = 6.50$, *** $p < 0.001$

**Fig 2C – Striatal TH immunofluorescence at 1, 2, and 4 m (in AU)**

| | AAV-mCh | | | AAV-hSyn | | | |
|---|---|---|---|---|---|---|---|
| | Mean | SEM | n | Mean | SEM | n | |
| 1m | 514.43 | 14.48 | 7 | 350.71 | 25.95 | 7 | *** |
| 2m | 506.14 | 37.37 | 7 | 376.57 | 33.15 | 7 | ** |
| 4m | 493.00 | 20.49 | 6 | 393.00 | 30.99 | 6 | * |

2 Way ANOVA
Main Effect of Time: $F(2,34) = 0.38$, $p = 0.6144$
Main Effect of Virus: $F(1,34) = 34.89$, ***$p < 0.0001$
Interaction: $F(2,34) = 0.49$, $p = 0.6144$

Post Hoc Tukey's Test:
1m: AAV-mCh < AAV-hSyn, *** $p = 0.0002$
2m: AAV-mCh < AAV-hSyn, ** $p = 0.0011$
4m: AAV-mCh < AAV-hSyn, * $p = 0.0158$

**Fig 2E – NeuN+ SNc cells at 1, 2, and 4 m (in # cells)**

| | AAV-mCh | | | AAV-hSyn | | | |
|---|---|---|---|---|---|---|---|
| | Mean | SEM | n | Mean | SEM | n | |
| 1m | 10948.50 | 381.35 | 6 | 10230.78 | 554.99 | 9 | n/a |
| 2m | 11307.50 | 293.39 | 6 | 9876.00 | 1306.86 | 5 | n/a |
| 4m | 10951.50 | 592.95 | 6 | 10849.43 | 1139.06 | 7 | n/a |

2 Way ANOVA
Main Effect of Time: $F(2,33) = 0.11$, $p = 0.9000$
Main Effect of Virus: $F(1,33) = 1.37$, $p = 0.2508$
Interaction: $F(1,33) = 1.37$, $p = 0.2508$

**Fig 2F – TH+ SNc cells, as percent of NeuN+ colabeled for TH at 1, 2, and 4 m**

| | AAV-mCh | | | AAV-hSyn | | | |
|---|---|---|---|---|---|---|---|
| | Mean | SEM | n | Mean | SEM | n | |
| 1m | 75.50 | 1.12 | 6 | 82.67 | 3.60 | 9 | n/a |
| 2m | 74.33 | 4.91 | 6 | 56.40 | 6.01 | 5 | n/a |
| 4m | 73.00 | 7.71 | 6 | 63.86 | 7.37 | 7 | n/a |

| | | | | | | | |
|---|---|---|---|---|---|---|---|
| | 2 Way ANOVA<br>Main Effect of Time: F(2,33) = 3.46, * $p$ = 0.0432<br>Main Effect of Virus: F(1,33) = 2.11, $p$ = 0.1553<br>Interaction: F(2,33) = 2.65, $p$ = 0.0855<br><br>Post Hoc Tukey's:<br>AAV-hSyn: 1m > 2m, * $p$ = 0.0223 | | | | | | |
| **Fig 4C (top) – [DA] release in dorsomedial striatum (FSCV)** | | | | | | | |
| | AAV-mCh | | | AAV-hSyn | | | |
| | Mean | SEM | n | Mean | SEM | n | |
| [DA] µM | 1.00 | 0.12 | 11 | 0.26 | 0.03 | 12 | *** |
| | Mann-Whitney U<br>U = 7, *** $p$ < 0.0001 | | | | | | |
| **Fig 4C (bottom) – [DA] release in dorsolateral striatum (FSCV)** | | | | | | | |
| | AAV-mCh | | | AAV-hSyn | | | |
| | Mean | SEM | n | Mean | SEM | n | |
| [DA] µM | 1.06 | 0.15 | 11 | 0.30 | 0.04 | 12 | *** |
| | Mann-Whitney U<br>U = 10, *** $p$ = 0.0002 | | | | | | |
| **Fig 4D – Striatal [DA] (HPLC)** | | | | | | | |
| | AAV-mCh | | | AAV-hSyn | | | |
| | Mean | SEM | n | Mean | SEM | n | |
| [DA] µM | 100.50 | 2.59 | 10 | 108.10 | 10.70 | 8 | ns |
| | Mann-Whitney U<br>U = 27, $p$ = 0.2743 | | | | | | |
| **Fig 5B – Whole cell electrophysiology, firing rate** | | | | | | | |
| | AAV-mCh | | | AAV-hSyn | | | |
| | Mean | SEM | n | Mean | SEM | n | |
| Firing Rate (Hz) | 2.35 | 0.23 | 22 | 2.87 | 0.38 | 19 | ns |
| | Unpaired t-test<br>t = 1.216, $p$ = 0.2311 | | | | | | |
| **Fig 5C – Whole cell electrophysiology, resting membrane potential (RMP)** | | | | | | | |
| | AAV-mCh | | | AAV-hSyn | | | |
| | Mean | SEM | n | Mean | SEM | n | |
| RMP (mV) | -52.21 | 0.79 | 22 | -51.90 | 0.97 | 19 | ns |
| | Unpaired t-test<br>t = 0.2512, $p$ = 0.8030 | | | | | | |
| **Fig 5D – Whole cell electrophysiology, Current/Frequency (reported in Hz)** | | | | | | | |
| | AAV-mCh | | | AAV-hSyn | | | |
| | Mean | SEM | n | Mean | SEM | n | |
| 0 pA | 2.824 | 0.422 | 17 | 2.50 | 0.34 | 16 | ns |
| 25 pA | 8.353 | 0.911 | 17 | 8.88 | 0.98 | 16 | |
| 50 pA | 10.235 | 0.838 | 17 | 15.63 | 1.71 | 16 | |
| 100 pA | 13.059 | 1.971 | 17 | 20.25 | 2.37 | 16 | |
| 125 pA | 13.647 | 1.656 | 17 | 24.50 | 2.83 | 16 | |
| 150 pA | 17.294 | 3.523 | 17 | 24.75 | 4.04 | 16 | |
| 175 pA | 16.118 | 2.933 | 17 | 27.38 | 4.21 | 16 | |
| | 2 Way ANOVA<br>Main Effect of Current: F (1.671, 51.81) = 31.84, ***$p$ < 0.0001<br>Main Effect of Virus: F (1, 31) = 5.827, *$p$ = 0.0219 | | | | | | |

| | | | | | | | |
|---|---|---|---|---|---|---|---|
| | Interaction: F(7, 217) = 3.889, ****p* < 0.0001<br>Repeated Measures: F (31, 217) = 10.76, ****p* < 0.0001<br><br>Bonferroni's Post Hoc Test:<br>100 pA: AAV-mCh < AAV-hSyn, **p* = 0.0233 | | | | | | |
| **Fig 5E – Whole cell electrophysiology, action potential half-width** | | | | | | | |
| | AAV-mCh | | | AAV-hSyn | | | |
| | Mean | SEM | n | Mean | SEM | n | |
| Half Width (msec) | 1.52 | 0.06 | 22 | 1.24 | 0.07 | 19 | ** |
| | Unpaired t-test<br>t = 0.2962, ** *p* = 0.0052 | | | | | | |
| **Fig 6B – Behavior, Velocity in open field test (reported in cm/sec) at 1, 2, and 4 m** | | | | | | | |
| | AAV-mCh | | | AAV-hSyn | | | |
| | Mean | SEM | n | Mean | SEM | n | |
| 1m | 2.48 | 0.11 | 31 | 2.22 | 0.10 | 31 | ns |
| 2m | 2.62 | 0.17 | 23 | 1.73 | 0.09 | 22 | *** |
| 4m | 2.47 | 0.14 | 12 | 2.14 | 0.16 | 15 | ns |
| | 2 Way ANOVA<br>Main Effect of Expression Time: F (2, 128) = 1.114, *p* = 0.3315<br>Main Effect of Virus: F (1, 128) = 19.43, *** *p* < 0.0001<br>Interaction: F (2, 128) = 3.738, * *p* = 0.0265<br><br>Bonferroni's Post Hoc Test:<br>2m: AAV-mCh < AAV-hSyn, *** *p* < 0.0001 | | | | | | |
| **Fig 6C – Behavior, Rotarod trial time (reported in sec) at 1, 2, and 4 m** | | | | | | | |
| | AAV-mCh | | | AAV-hSyn | | | |
| | Mean | SEM | n | Mean | SEM | n | |
| 1m | 113.06 | 5.30 | 31 | 123.00 | 4.47 | 31 | n/a |
| 2m | 127.75 | 6.91 | 24 | 117.50 | 5.19 | 22 | n/a |
| 4m | 135.67 | 12.66 | 12 | 127.13 | 4.29 | 15 | n/a |
| | 2 Way ANOVA<br>Main Effect of Expression Time: F (2, 129) = 1.988, *p* = 0.1412<br>Main Effect of Virus: F (1, 129) = 0.3087, *p* = 0.5794<br>Interaction: F (2, 129) = 1.908, *p* = 0.1525 | | | | | | |
| **Fig 6D – Behavior, Pole test trial time (reported in sec) at 1, 2, and 4 m** | | | | | | | |
| | AAV-mCh | | | AAV-hSyn | | | |
| | Mean | SEM | n | Mean | SEM | n | |
| 1m | 11.25 | 0.83 | 22 | 20.00 | 7.56 | 27 | n/a |
| 2m | 13.93 | 3.18 | 24 | 20.43 | 3.79 | 21 | n/a |
| 4m | 8.86 | 1.30 | 12 | 12.68 | 2.13 | 15 | n/a |
| | 2 Way ANOVA<br>Main Effect of Expression Time: F (2, 115) = 0.7637, *p* = 0.4683<br>Main Effect of Virus: F (1, 115) = 2.440, *p* = 0.1210<br>Interaction: F (2, 115) = 0.1147, *p* = 0.1210 | | | | | | |
| **Fig Supp 1A (left) – Cell counts in SNc, AAV-mCh v hSyn (1:5 dilution)** | | | | | | | |
| | AAV-mCh | | | AAV-hSyn | | | |
| | Mean | SEM | n | Mean | SEM | n | |
| NeuN+ (# cells) | 11764.25 | 893.45 | 4 | 10866.25 | 1142.21 | 4 | ns |
| TH+ (# cells) | 9111.75 | 809.30 | 4 | 8027.00 | 887.48 | 4 | ns |

| | | | | | | | |
|---|---|---|---|---|---|---|---|
| mCh+ (# cells) | 5057.50 | 217.50 | 4 | 5259.50 | 638.84 | 4 | ns |
| hSyn+ (# cells) | 0.00 | 0.00 | 4 | 6402.75 | 756.96 | 4 | *** |
| | 2 Way ANOVA<br>Main Effect of Cellular Marker: $F(3,24) = 45.35$, ***$p < 0.0001$<br>Main Effect of Virus: $F(1,24) = 4.671$, * $p = 0.0409$<br>Interaction: $F(3,24) = 10.98$, ***$p < 0.0001$<br><br>Bonferroni's Post Hoc Test:<br>hSyn+: AAV-mCh < AAV-hSyn, ***$p < 0.0001$ | | | | | | |
| **Fig Supp 1A (center) – NeuN+ SNc cells at 1, 2, and 4 m (in # cells) (1:5 dilution)** | | | | | | | |
| | AAV-mCh | | | AAV-hSyn | | | |
| | Mean | SEM | n | Mean | SEM | n | |
| 1m | 11764.25 | 893.45 | 4 | 10866.25 | 1142.21 | 4 | n/a |
| 2m | 11251.00 | 679.47 | 4 | 10676.50 | 203.90 | 4 | n/a |
| 4m | 12834.75 | 646.52 | 4 | 12257.00 | 792.01 | 4 | n/a |
| | 2 Way ANOVA<br>Main Effect of Time: $F(2,18) = 2.268$, $p = 0.1323$<br>Main Effect of Virus: $F(1,18) = 1.151$ $p = 0.2975$<br>Interaction: $F(2,18) = 0.028$, $p = 0.9721$ | | | | | | |
| **Fig Supp 1A (right) – TH+ SNc cells (% NeuN+ colabeled for TH) at 1, 2, and 4 m (1:5 dilution)** | | | | | | | |
| | AAV-mCh | | | AAV-hSyn | | | |
| | Mean | SEM | n | Mean | SEM | n | |
| 1m | 77.00 | 2.80 | 4 | 74.00 | 4.24 | 4 | n/a |
| 2m | 70.50 | 3.59 | 4 | 65.75 | 2.21 | 4 | n/a |
| 4m | 72.00 | 5.37 | 4 | 67.50 | 2.40 | 4 | n/a |
| | 2 Way ANOVA<br>Main Effect of Time: $F(2,18) = 2.393$, $p = 0.1286$<br>Main Effect of Virus: $F(1,18) = 1.918$, $p = 0.1830$<br>Interaction: $F(2,18) = 0.034$, $p = 0.9663$ | | | | | | |
| **Fig Supp 1B – Striatal [DA] (HPLC)** | | | | | | | |
| | AAV-mCh | | | AAV-hSyn | | | |
| | Mean | SEM | n | Mean | SEM | n | |
| [DA] µM | 116.90 | 5.09 | 10 | 142.8 | 7.13 | 11 | * |
| | Unpaired t-test<br>$t = 2.841$, $p = 0.0101$ | | | | | | |
| **Fig Supp 1C (left) – [DA] release in dorsomedial striatum (FSCV) (1:5 dilution)** | | | | | | | |
| | AAV-mCh | | | AAV-hSyn | | | |
| | Mean | SEM | n | Mean | SEM | n | |
| [DA] µM | 1.630 | 0.1639 | 14 | 0.5993 | 0.1559 | 14 | *** |
| | Unpaired t-test<br>$t = 4.557$, $p = 0.0001$ | | | | | | |
| **Fig Supp 1C (right) – [DA] release in dorsolateral striatum (FSCV) (1:5 dilution)** | | | | | | | |
| | AAV-mCh | | | AAV-hSyn | | | |
| | Mean | SEM | n | Mean | SEM | n | |
| [DA] µM | 1.442 | 0.1656 | 14 | 0.7700 | 0.1717 | 14 | ** |
| | Unpaired t-test<br>$t = 2.818$, $p = 0.0091$ | | | | | | |
| **Fig Supp 1D – Behavior, Velocity in open field test (reported in cm/sec) at 1, 2, and 4 m (1:5 dilution)** | | | | | | | |
| | AAV-mCh | | | AAV-hSyn | | | |

| | | | | | | | |
|---|---|---|---|---|---|---|---|
| | Mean | SEM | n | Mean | SEM | n | |
| 1m | 2.27 | 0.12 | 13 | 2.02 | 0.11 | 17 | ns |
| 2m | 2.28 | 0.15 | 8 | 1.80 | 0.08 | 12 | * |
| 4m | 2.31 | 0.23 | 6 | 1.76 | 0.12 | 7 | ns |
| | 2 Way ANOVA<br>Main Effect of Expression Time: F (2, 57) = 0.4913, *p* = 6144<br>Main Effect of Virus: F (1, 57) = 14.58, *** *p* = 0.0003<br>Interaction: F (2, 57) = 0.7661, * *p* = 0.4695<br><br>Bonferroni's Post Hoc Test:<br>2m: AAV-mCh < AAV-hSyn, * *p* = 0.0624 | | | | | | |
| **Fig Supp 1E – Behavior, Rotarod trial time (reported in sec) at 1, 2, and 4 m (1:5 dilution)** | | | | | | | |
| | AAV-mCh | | | AAV-hSyn | | | |
| | Mean | SEM | n | Mean | SEM | n | |
| 1m | 131.54 | 9.68 | 13 | 133.82 | 9.08 | 17 | n/a |
| 2m | 137.33 | 9.92 | 9 | 115.62 | 8.48 | 13 | n/a |
| 4m | 126.83 | 8.20 | 6 | 130.29 | 10.13 | 7 | n/a |
| | 2 Way ANOVA<br>Main Effect of Expression Time: F (2, 59) = 0.2389, *p* = 0.7882<br>Main Effect of Virus: F (1, 59) = 0.3856, *p* = 0.5370<br>Interaction: F (2, 59) = 1.006 *p* = 0.3718 | | | | | | |
| **Fig Supp 1F – Behavior, Pole test trial time (reported in sec) at 1, 2, and 4 m (1:5 dilution)** | | | | | | | |
| | AAV-mCh | | | AAV-hSyn | | | |
| | Mean | SEM | n | Mean | SEM | n | |
| 1m | 13.54 | 2.24 | 10 | 11.12 | 1.80 | 12 | n/a |
| 2m | 12.15 | 2.28 | 9 | 11.13 | 1.73 | 13 | n/a |
| 4m | 9.20 | 2.06 | 5 | 7.00 | 0.38 | 7 | n/a |
| | 2 Way ANOVA<br>Main Effect of Expression Time: F (2, 50) = 1.991, *p* = 0.1473<br>Main Effect of Virus: F (1, 50) = 1.233, *p* = 0.2721<br>Interaction: F (2, 50) = 0.0801, *p* = 0.9231 | | | | | | |

**Supplementary Table 1 (associated with Figures 1-6).** Statistical Table

| | AAV-mCh (n = 22) | AAV-hSyn (n = 19) |
|---|---|---|
| Basal Activity | | |
| Basal Firing Rate | 2.35 ± 0.23 Hz | 2.87 ± 0.38 Hz |
| Coefficient of Variation | 21.64 ± 5.08 | 19.70 ± 5.53 |
| Resting Membrane Potential | -52.21 ± 0.79 mV | -51.90 ± 0.97 mV |
| Action Potential Measures | AAV-mCh (n = 22) | AAV-hSyn (n = 19) |
| Half Width | 1.52 ± 0.06 ms | 1.24 ± 0.07 ms** |
| AHP | -60.30 ± 1.06 mV | -59.72 ± 1.40 mV |
| Upstroke | 107.9 ± 8.34 V/s | 163.8 ± 13.87 V/s*** |
| Downstroke | -61.49 ± 2.80 V/s | -76.48 ± 5.37* |

Table 2. Intrinsic properties of midbrain dopamine neurons at 1 month post AAV injection. All data shown as mean ± SEM. All comparisons were unpaired t-tests between AAV-mCh and AAV-hSyn, * $p < 0.05$, ** $p < 0.01$, *** $p < 0.001$.

**Supplementary Table 2 (associated with Figure 5).** DAT-Cre mice were injected in the substantia nigra pars compacta (SNc) with AAV encoding mCherry or human wild-type ⟨-synuclein and mCherry, and sacrificed after 1 month for *ex vivo* patch-clamp recordings of SNc dopamine neurons. Intrinsic properties were determined in whole-cell current-clamp mode.

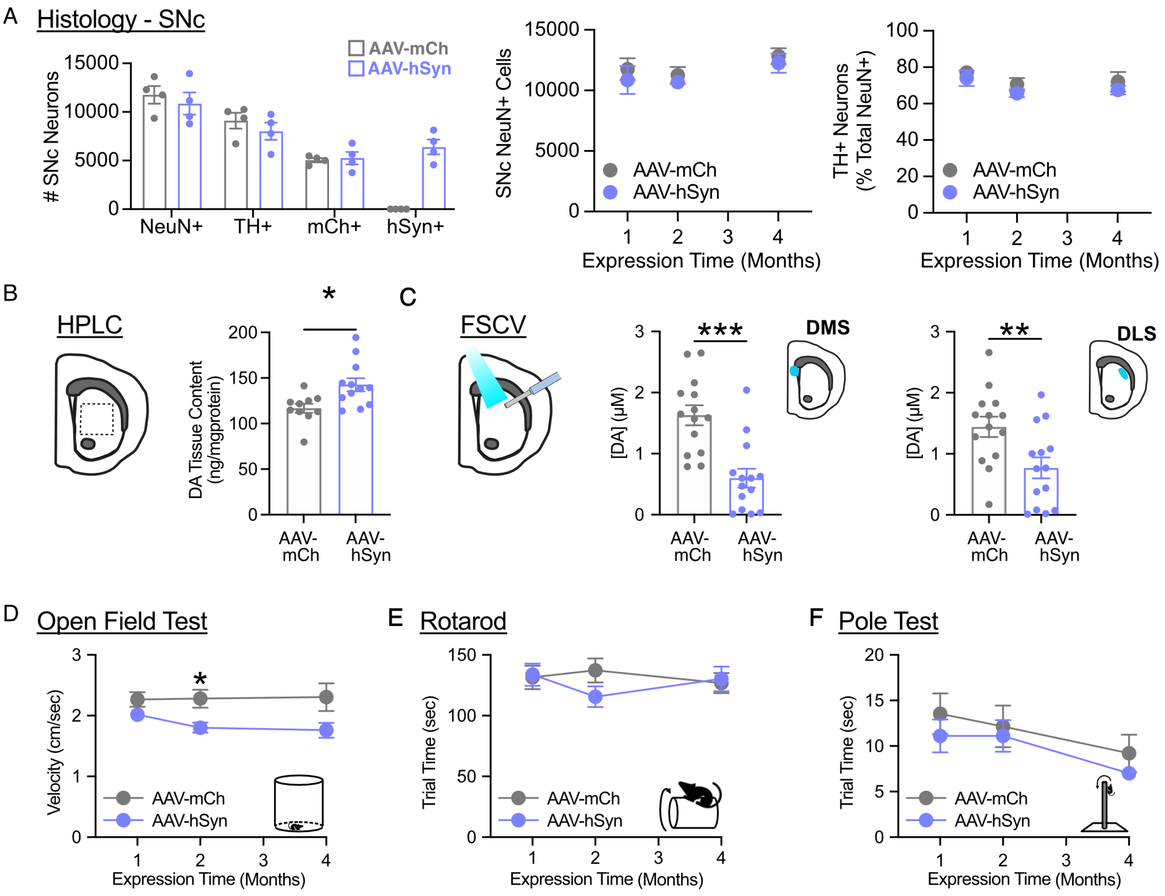

**Supplementary Figure 1: Dilution of the AAV encoding α-synuclein leads to more modest pathological, physiological, and motor defects.** DAT-Cre mice were injected in the substantia nigra pars compacta (SNc) with AAV encoding mCherry or human wild-type α-synuclein and mCherry. AAV was diluted at 1:5 (1 uL of AAV to 4 uL of sterile saline) prior to injection. (A) Animals were sacrificed at 1, 2, or 4 months for postmortem histology. Left: Summary of SNc neurons expressing NeuN, TH, mCherry, and human ⟨-synuclein in each group. Middle: Summary of the total number of neurons (NeuN-positive) in the SNc. There was no significant change between groups or across timepoints in animals injected with diluted AAVs ($p = 0.1830$). Right: Summary of TH+ neurons, as measured by percent of NeuN+ cells co-labeled for TH, which was unchanged in mice injected with diluted AAV-hSyn. (B) Animals were sacrificed at 1 month for HPLC assessment of dopamine content in striatal tissue punches. Dopamine content was not reduced in animals treated with diluted AAV-hSyn and, on the contrary, these animals displayed a small but significant increase ($p = 0.0101$). (C) Animals were sacrificed at 1 month for ex vivo fast-scan cyclic voltammetry. Summary of optically evoked dopamine release in slices from mice treated with diluted AAV-mCh or AAV-hSyn in the DMS (left) and DLS (right). Dopamine release was reduced in both regions ($p = 0.0001$; $p = 0.0091$) (D-F) Mice injected with diluted AAV-mCh and AAV-hSyn were tested at 1, 2, and 4 months on the open field test (D), accelerating rotarod (E) and pole test (F). There was a reduction in open field locomotor velocity at 2 months in the AAV-hSyn group ($p = 0.0003$, Post Hoc Bonferroni's Test, $p = 0.0624$), but no change in any other motor assays (diluted AAV-mCh: N=13, 8-9, 6; diluted AAV-hSyn: N = 17, 12-13, 7). For pole test, n was lower at the 1 month time point, at 10 and 12, respectively, for AAV-mCh and AAV-hSyn. All data shown as mean +/- SEM. Superimposed dots represent individual mice.

### Methods

#### Mice

DAT-IRES-Cre mice on a C57Bl/6 background (RRID:IMSR_JAX:006660) were crossed to wild-type C57Bl/6 mice (JAX) to yield male and female hemizygous DAT-Cre mice for use in experiments. In a subset of experiments that measured evoked dopamine release, DAT-Cre mice were crossed to the Cre-dependent Channelrhodopsin-2 line Ai32 (RRID:IMSR_JAX:012569). Animals were housed 1-5 per cage on a 12-hour light/dark cycle with *ad libitum* access to rodent chow and water. All behavioral assays were performed during the light phase. We complied with local and national ethical and legal regulations regarding the use of mice in research. All experimental protocols were approved by the University of California, San Francisco, University of Colorado Anschutz Medical Campus, and National Institutes of Health Institutional Animal Care and Use Committees.

#### Surgery

A detailed protocol for stereotaxic surgery can be found at dx.doi.org/10.17504/protocols.io.n2bvj6qynlk5/v1.

Briefly, surgeries were performed at 2-4 months of age. Anesthesia was induced with intraperitoneal (IP) injection ketamine/xylazine and maintained with 0.5%-1.0% inhaled isoflurane. Mice were placed in a stereotaxic frame, and the head was adjusted until the top surface of the skull was flat. To achieve a flat skull surface, the tip of a drill bit in a stereotax-mounted drill was placed at lambda and bregma, with the head and ear bars adjusted until there was <0.1 mm of vertical difference between the two sites. Likewise, spots 2 mm to the left and right of bregma were used to achieve a flat surface from side to side (0 mm vertical difference between the two sites). The drill was then used to create holes over the bilateral substantia nigra pars compacta (SNc). AAV5-DIO-IRES-mCherry or AAV5-DIO-⟨syn-IRES-mCherry (both 300 nL, Vector Builder, titer: $10^{13}$ particles/⌠L, undiluted) was injected in each side of the SNc (-2.1 AP, +/-1.2 ML, -4.5 DV). The injection needle was left in place for 10 minutes prior to being withdrawn and the scalp being sutured. Animals were then administered buprenorphine (IP) and ketoprofen or carprofen (SQ) for postoperative analgesia, and monitored closely over the subsequent week for full recovery. To ensure consistency of targeting across laboratories, we performed cross-validation experiments by injecting either dye or the virus itself, and performing postmortem histology (see below). Methodology was iterated until therefore was relative consistency both within and between sites.

**Postmortem Histology**

Detailed protocols for brain slicing, immunofluorescence staining, and imaging can be found here:
dx.doi.org/10.17504/protocols.io.14egn7yyzv5d/v1
https://dx.doi.org/10.17504/protocols.io.dm6gp3mo5vzp/v1
https://dx.doi.org/10.17504/protocols.io.n92ldm1nnl5b/v2
https://dx.doi.org/10.17504/protocols.io.rm7vzk2k8vx1/v1

Briefly, mice were sacrificed at 1, 2, and 4 months after AAV injections. Mice were deeply anesthetized with ketamine-xylazine (IP) or inhaled isoflurane and transcardially perfused with 4% paraformaldehyde (PFA) in PBS. The brain was then dissected and placed in 4% PFA overnight. The brain was then transferred to a solution of 30% sucrose in PBS for storage. All brains used for semi-quantitative and quantitative histology were shipped to a central laboratory (University of Sydney) for subsequent analysis.

Briefly, thin sections were prepared on a sliding microtome Thermo Fisher Epredia™ CryoStar™ NX50 (Thermo Fisher Scientific, North Ryde, Australia). Coronal sections containing the striatum (20 microns) were cut from Bregma 1.69mm to -0.95mm. Coronal sections containing the SNc (30 microns) were cut from Bregma -2.69mm to Bregma -4.03mm (anatomical location as referenced (Paxinos, 2004)). Consecutive coronal sections were collected into eight tubes (on average twelve striatal sections per tube and three midbrain sections per tube) prior to freezing in cryoprotectant (30% sucrose, 30% glycerol in 0. M Phosphate buffer) for storage at -20°C to preserve tissue integrity prior to use.

Sections were then processed for immunohistochemistry. These experiments used triple-labelled immunofluorescence to identify neuronal structures expressing and not expressing human α-synuclein. Sections were first treated with citrate buffer (pH 6.0) at 85°C for 30 mins, followed by blocking buffer (0.3% Triton X-100 and 5% horse serum in 1 x Tris buffered saline) for 1 hour. Sections were incubated in the cocktail of primary antibodies (Table 1) on a shaker at 50 rpm at 4°C for two days followed by three washes with 1 x Tris buffered saline before incubation in the cocktail of secondary antibodies (Table 1) containing Hoechst (1:2000, 34580, North Ryde, Australia) for 2 hrs at room temperature. Additionally, neuronal pathology was revealed either by

Thioflavin-S (0.05%, T1892, Sigma Aldrich, Melbourne, Australia) staining for 10 mins at room temperature to detect fibrilized α-synuclein (Stojkovska & Mazzulli, 2021) or with immunofluorescence labelling using pathological α-synuclein and p62 antibodies (Table 1). Sections were washed, mounted on slides, and coverslipped with anti-fade fluorescent mounting medium (Dako, Santa Clara, United States). Negative controls were included for each staining batch by omitting primary or secondary antibodies as well as using *SNCA* knockout mouse brain tissue.

All images for quantification were photographed at 20x magnification using an Olympus VS200 slide scanner equipped with X-line high-performance objectives to avoid stitching artifacts for downstream analysis (Olympus, Tokyo, Japan). Serial sections were aligned according to the midline and ventral bottom of individual image and corrected for possible displacement and distortion by moving and/or rotating them individually as necessary based on their original stereological sequence prior to importing into the AI program QuPath version 0.4.3 (RRID:SCR_018257; Bankhead et al., 2017) for quantitation. The location of TH immunoreactivity and the mouse brain atlas in stereotaxic coordinates (Paxinos, 2004) were used to identify standard landmarks and select three of 12 stained serial sections of the striatum (between Bregma 1.93 to -0.95 mm) and all six stained serial sections of the midbrain for quantitation. The midbrain rostral, intermediate and caudal sections were between Bregma -2.91 to -3.15mm, -3.15 to -3.39mm, and -3.39 to -3.63mm respectively.

For descriptive analyses, images were photographed at 40x magnification using a confocal microscope (Nikon C2, Tokyo, Japan) and 62x magnification using an Olympus VS200 slide scanner. Image deconvolution was processed with Olympus TruSight deconvolution software (Olympus, Tokyo, Japan) to flatten the stained structures into a single plane.

***Ex vivo* Fast-Scan Cyclic Voltammetry**

A detailed protocol for preparation of brain slices and performing fast-scan cyclic voltammetry can be found at https://www.protocols.io/view/fast-scan-cyclic-voltammetry-fscv-in-mouse-brain-s-cyjjxukn.

Briefly, animals were deeply anesthetized, transcardially perfused, decapitated, and the brain dissected. 240 μm brain slices containing the striatum were prepared and transferred to a 34° C holding chamber containing artificial cerebrospinal fluid (ACSF) bubbled with 5% $CO_2$ in $O_2$ and containing in mM: 126 NaCl, 2.5 KCl, 1.2 $MgCl_2$, 1.2 $NaH_2PO_4$, 2.5 $CaCl_2$, 21.4 $NaHCO_3$, and

11.1 D-glucose. This ACSF contained 10 µM MK-801 and 34° C and slices were held for 45 minutes or more prior to voltammetry.

Brain slices were placed into a recording chamber perfused with warm (32°C), carbogenated ACSF containing in mM 1 uM dihydro-beta-erthroidine (Tocris, CAS: 29734-68-7) and 0.5 uM sulpiride (Tocris, CAS: 23672-07-3). A carbon-fiber recording electrode was placed into the striatum for measurement of evoked dopamine release. After a period of calibration and stabilization (approximately 5 stimuli), recordings were performed using single 1 msec blue light pulses (473 nm, 5 mW). The average of the subsequent 3 sweeps was calculated to determine dopamine release.

Assessment of Dopamine Concentration by High Performance Liquid Chromatography (HPLC)

A detailed protocol for the collection of striatal slices for HPLC can be found at dx.doi.org/10.17504/protocols.io.3byl4qj4zvo5/v1.

Briefly, animals were deeply anesthetized and decapitated. Unlike other experiments in this study, these animals were not transcardially perfused. The brain was transferred immediately to cold Hank's Balanced Salt Solution (HBSS) containing 10 mM HEPES and 20 mM glucose and 1 mm coronal slices were collected using a rodent brain matrix. From the slices containing the center of the striatum, the striatum was isolated and flash frozen in liquid nitrogen. Slices were stored at -80C until use. The tissues were sent on dry ice to the Neurochemistry Core at the Vanderbilt University Medical Center, where they measured total tissue protein and tissue dopamine using HPLC.

***Ex vivo* Electrophysiology**

A detailed protocol for preparation of midbrain slices and whole-cell patch-clamp recordings can be found at dx.doi.org/10.17504/protocols.io.kxygx7km4l8j/v1.

Briefly, control (IRES-mCherry) or ⟨syn (⟨syn-IRES-mCherry) mice were deeply anesthetized with inhaled isoflurane, transcardially perfused, decapitated, and the brain dissected. 200 um coronal brain slices containing the midbrain were prepared and transferred to a warmed ACSF holding chamber for 45 minutes or more. Slices were then transferred to an electrophysiology setup, where they were superfused with ACSF.

Within the SNc, cells expressing mCherry were targeted for whole-cell patch-clamp recordings with a potassium-based physiological internal solution. For each cell, spontaneous action potential firing, resting membrane potential, input resistance, and responses to brief (1.5 second) current injections were recorded in current-clamp mode. The average firing rate is calculated as the total number of spikes divided by the total time in sweeps of spontaneous firing. Evoked firing rate was calculated as the total number of spikes during the 1.5 second current injection, divided by 1.5 seconds. The resting membrane potential was calculated as the average membrane potential during the interspike interval in sweeps of spontaneous firing. The action potential half-width and other aspects of spike shape are calculated during periods of spontaneous firing. The half-width is the width of the action potential at half of the spike height (action potential peak – threshold). The input resistance was calculated from the voltage deflection in response to small negative current steps.

**Behavioral Assays**

A detailed protocol for all of the motor assays used here can be found at https://www.protocols.io/view/motor-behavior-assays-mouse-q26g7yo4kgwz/v1.

Briefly, mice were tested during the light phase with three different motor assays: the open field test, the accelerating rotarod, and the pole test. Control and ⟨syn mice were included in each behavioral cohort, and the experimenter was blinded to each animal's group assignment (control or ⟨syn).

For the open field test, animals were habituated to the chamber (a transparent acrylic cylinder) for two days (30 minutes each) prior to the first testing session, and then tested (20 minutes) at up to three time points: 1, 2, and 4 months after AAV injection. For the pole test, animals were trained for two days prior to the test session (3 trials each), and then tested on the third day (3 trials). In each trial the animal was placed at the top of the pole, facing up. The time to turn (facing down), the time to descend to the ground (two front paws on the ground), and total time were recorded.

For the accelerating rotarod test, animals were tested on a single day (3 trials). Animals were placed on the accelerating rotarod (0-80 rpm) and monitored until they fell off the rotarod, or held on for a full rotation. The fall time was recorded for each trial.

**Quantification and Statistical Analysis**

The details of statistical comparisons, including the associated figure panels, comparison groups, N, statistical tests, and p values can be found in the Supplementary Table 1 (Statistical Table). For physiological assays (*ex vivo* electrophysiology and fast-scan cyclic voltammetry), animals were excluded if there was not strong mCherry expression in the SNc, as assayed in living slices under the microscope. For behavioral assays, animals were excluded based on blinded semi-quantitative assessment of mCherry expression by postmortem histology. Details of this approach can be found at dx.doi.org/10.17504/protocols.io.81wgbrmknlpk/v1.

To assess the impact of control and hSyn AAVs on striatal TH-immunoreactive terminals, the striatal borders were annotated in QuPath using the outline tool based on the location of the TH-immunoreactivity. Four striatal subregions (dorsomedial, dorsolateral, ventromedial, and ventrolateral quadrants) were separated by drawing arbitrary vertical and horizontal lines crossing the striatal center. For each subregion, the mean fluorescence intensity was measured by the computed intensity algorithm (parameters: intensity features of 2 μm pixel size and 25 μm tile diameter) normalised to the background signal measured from the cingulum in each section. Both quadrant and entire section values were exported, and the averaged value (from a minimum of three striatal sections per mouse) was used for the statistical analysis.

To assess the impact of control and hSyn AAVs on midbrain TH-immunoreactive neurons, the compact region of the SNC was separated from the remaining ventral tegmental region (VTA) in each section using the outline tool in QuPath and all neurons (NeuN+), dopamine neurons (TH+), AAV-impacted neurons (mCherry+), and AAV- hαSyn transfected neurons (15G7+/BD-1+) were quantified. The algorithm for neuronal number was based on the cell detection function with objective classification for positive neurons in the targeted channel, followed by a single measurement classifier. The estimated true number (N) of neurons within each region in each section from the QuPath single objective classifier was -

$$N = n \times \left(\frac{t}{t+H}\right)$$

n is the counted number, t is the mean section thickness (30μm), and H is the mean height of the neurons (8.324μm). The total neuron number for each mouse was further calculated by multiplying the estimated number of counted neurons in the six spaced serial sections by the number of series of sections used (8/2 series used).
